# Transposable element activity in the transcriptomic analysis of mouse pancreatic tumors

**DOI:** 10.1101/2021.07.16.452652

**Authors:** Emmanuelle Lerat, Nelly Burlet, Vincent Navratil, Camilles Noûs

**Affiliations:** Université de Lyon, Université Lyon 1, CNRS, Laboratoire de Biométrie et Biologie Evolutive UMR 5558, F-69622 Villeurbanne, France; PRABI, Rhône Alpes Bioinformatics Center, UCBL, Université Claude Bernard Lyon 1, F-69000 Lyon, France; Laboratoire Cogitamus, https://www.cogitamus.fr/

## Abstract

Transposable elements (TEs) are middle-repeated DNA sequences that can move along chromosomes using internal coding and regulatory regions. By their ability to move and because they are repeated, TEs can promote mutations. Especially they can alter the expression pattern of neighboring genes and have been shown to be involved in the mammalian regulatory network evolution. Human and mouse share more than 95% of their genomes and are affected by comparable diseases, which makes the mouse a perfect model in cancer research. However not much investigation concerning the mouse TE content has been made on this topics. In human cancer condition, a global activation of TEs can been observed which may ask the question of their impact on neighboring gene functioning. In this work, we used RNA sequences of highly aggressive pancreatic tumors from mouse to analyze the gene and TE deregulation happening in this condition compared to pancreas from healthy animals. Our results show that several TE families are deregulated and that the presence of TEs is associated with the expression divergence of genes in the tumor condition. These results illustrate the potential role of TEs in the global deregulation at work in the cancer cells.

## Introduction

Mammalian genomes contain a high proportion of repeated sequences when compared to that of the protein coding genes. For example, the human genome harbors only about 2% of coding genes whereas transposable elements (TEs) make up 45% of this genome (1). These values are roughly similar in the mouse genome even if the proportion of TEs is slightly inferior with 38.55% (2). TEs are middle-repeated DNA sequences that have the ability to move from one position to another along chromosomes. They typically encode for all the proteins necessary for their movement and possess internal regulatory regions, allowing their independent expression. Different categories of TEs have been identified according to their transposition intermediates and their structural features (3). Two main classes have been described. The first comprises the retrotransposons, which use an RNA intermediate and in which are found the LTR-retrotransposons (endogenous retrovirus-like mobile elements) and the non-LTR retrotransposons grouping the LINE and the SINE elements (standing for Long and Short Interspersed Nuclear Elements respectively). The second class groups the DNA transposons, which use a DNA intermediate. Although the global proportion of TEs between the mouse and the human genomes are similar, there are some notable differences. In particular, lineage-specific TEs are more numerous in the mouse than in the human genome, indicating a more recent activity (2). This activity can be ascertained when observing different mouse strains for which several thousand of insertion variants were detected, although only a small fraction were shown to promote changes in nearby gene expression (4). As in human, the mouse genome harbors a majority of non-LTR retrotransposons (27.42% corresponding to 19.20% of LINEs and 8.22% of SINEs), followed by LTR retrotransposons (9.87%) and very few DNA transposons (0.88%) (2). However, variations exist concerning the different proportions of specific families inside each class. For example, ERV-2 LTR retrotransposons are more abundant in mouse than in human. Moreover, LTR retrotransposons present several active families in mouse whereas only one is active in the human genome (5). Despite these differences, TE insertions have been detected in orthologous locations between both genomes (2).

By their ability to move and because they are repeated, TEs can promote various types of mutations, which are expected to be mostly deleterious when affecting functional regions. Indeed, when TE insertions occur in or near protein-coding genes, they can lead to the disruption of coding capacity or to the alteration of the splicing or of the polyadenylation patterns (6). Moreover, because TEs possess their own regulatory sequences, they can alter the expression pattern of neighboring genes (6). Finally, the possibility of homologous recombination between copies can promote illegitimate recombinations, chromosome breakages, deletions and genome rearrangements (6). In human, 0.3 % of TE insertions have been suggested to be responsible for diseases (7) and approximately 96 new transposition events have been directly linked to single-gene diseases (8). For example, the Alport syndrome has been shown to be due to a TE mediated rearrangement resulting in the partial deletion and fusion of two genes (9). An Alu insertion localized inside an exon of the ClC-5 gene has been proposed to cause exon skipping by interfering with splicing regulatory elements, which lead to the Dent’s disease (10). These examples show evidences of the direct implication of TE insertions in the induction of diseases. However, mammalian genomes harbor millions of fixed TE insertions that have not been counter-selected during evolution or because they were on the contrary selected for some advantages they provided to the organism. Recently, it has been shown that the co-localization of elements from old repeat families may explain the global conservation of genome folding observed between homologous regions of the human and mouse genomes, which would indicate a contribution of these elements in maintaining and/or reshaping genome architecture over evolutionary times (11). The impact of the TEs on the regulatory network evolution has also been observed. For example, a comprehensive survey of transcription factor binding sites in human and mouse showed that 20% of these sites were embedded within TEs with, on average, 66% of them being cell type-specific (12). These results suggest that TEs have shaped gene regulatory networks during mammalian evolution. Moreover, some TE functional impact on mammalian developmental processes have been described. For instance, a TE-derived promoter generates a cell type-specific isoform of the mouse gene Dicer1, which is essential for oogenesis (13). Similarly, many other genes in mouse oocytes and zygotes are expressed from endogenous retrovirus insertions, which act as alternative promoters (14). The investigation of LTR retrotransposons suspected to enhance the activity of host genes showed that some of them are essential for normal gene expression in early mouse development (15). Therefore, fixed TE insertions could still influence the genome evolution, expression or regulation (16).

In cancer condition, in addition of being sometimes the causative agent like for example in a colorectal cancer (17), global activation of TEs has often been observed along with epigenetic changes (18). This is the case, for example, of Alu elements in a human brain tumor, which display a loss of methylation (19). More globally, one of the common characteristics to all cancer types is a general loss of methylation of all TEs (20). Consequently, particular endogenous retroviruses in human produce viral particles in melanoma cells (21), TE expression is enhanced in urothelial and renal carcinoma cells (22), in some carcinomas (23), in human leukemia (24), and in human colorectal, ovarian and breast cancers (25–28). The TE reactivation can have an impact on neighboring genes. For example, in a previous work, we have shown that the presence of SINEs is associated with the expression divergence of neighboring genes in human cancer cells, with an increasing impact as the number of SINEs near genes increases (29). The analysis of TE-derived oncogene transcripts from 15 cancer types allowed to show that the onco-exaptation, the mechanism by which cryptic regulatory elements within TEs can be epigenetically reactivated in cancer to influence oncogenesis, could be a prevalent mechanism contributing to the oncogene activation (30).

Since human and mice share more than 95% of their genomes and are affected by comparable diseases, the results of mouse experiments most of the time relate to human biology, which makes the mouse a perfect model in cancer research. However not much investigation concerning the mouse TE content on this topics has been made. In this work, we used RNA-seq data obtained from genetically engineered mice (31) presenting highly aggressive pancreatic tumors to analyze the gene and TE deregulation happening in this condition compared to pancreas from healthy animals. Our results show that the presence of TEs is clearly associated with the expression divergence of genes in the tumor condition, which illustrate their potential role in the global deregulation at work in the cancer cells.

## Material and Methods

### Ethical statement

The experiments were performed in compliance with the animal welfare guidelines of the European Union, the French legislation and the ARRIVE guidelines. The experiments were approved by the CECCAPP Région Rhône-Alpes ethical committee https://www.sfr-biosciences.fr/ethique/ceccapp/ceccapp under the control of the Ministère de l’Enseignement Supérieur et de la Recherche (#C2EA15) http://www.enseignementsup-recherche.gouv.fr/cid70597/l-utilisation-des-animaux-a-des-fins-scientifiques.html.

### Sample collection and sequencing of Illumina RNA libraries

We obtained two replicates from the shredding and lysis of pancreatic tumor tissues (one quarter of pancreas maximum) corresponding to pancreatic ductal adenocarcinoma (PDAC) from Pdx1-Cre; LSL-KrasG12D; Ink4a/Arflox/lox mice in Guanidinium Thiocyanate Buffer (Guanidine Thiocyanate 5M, Sodium Citrate 0.5M pH7, Lauryl Sarcosine 10%, B-mercaptoethanol 1%). Extractions of RNA using Qiagen RNeasy Plus Mini kit were performed according to the manufacturer instructions. After DNAses treatment, the quantity and the integrity (RIN measurement) of RNA were measured using Bioanalyser 2100 (Agilent). Two replicates of normal pancreas Balb/C mouse were obtained from Clinisciences for which ontamination by RNase, genomic DNA polysaccharides, and proteoglycans has been eliminated.

PolyA+ mRNAs were sequenced by Illumina Technology in HiSeq 3000 sequencers from the MGX-Montpellier GenomiX facility (https://www.mgx.cnrs.fr/ ; Montpellier, France). The four samples were sequenced in 2×150 bp reads with an average insert size of 350 bp and produced about 100 Millions of reads per sample. All data have been deposited on the SRA website (project ID PRJNA691737). The sequencing quality of each sample was evaluated using the FastQC software version 0.11.2 (32). Although the data were globally of very good quality, the reads were cleaned using the UrQt software (33) with parameter t=10 to remove low quality bases.

### Differential expression analysis of genes

To identify differentially expressed genes between cancer and normal tissues, we followed the protocol proposed by Pertea et al. (34). The reads were mapped using Hisat2 (35) on the mouse reference genome version GRCm38.p6 downloaded on the NCBI website (https://www.ncbi.nlm.nih.gov/assembly/GCF_000001635.26/). For all four samples, the overall alignment rate was about 94%. Transcripts were then assembled and quantify from the mapped reads using Stringtie (36) and the annotated gtf file from the mouse reference genome (Mus_musculus.GRCm38.97.gtf). The differential expression analysis was performed on the read count produced by Stringtie using DESeq2 (37). It has to be noted that the analyses were performed using two replicates for each sample, which could prevent the identification of differentially expressed genes with very low fold changes and could also lead to some false positive detection32. To prevent this issue we used a very stringent p-value threshold. In that respect, transcripts are classified as significantly differentially expressed if the p-value, after correction for multiple tests with the False Discovery Rate (FDR), is below 0.01. To determine the number of genes differentially expressed between the two conditions, we retrieved the Ensembl ID of genes corresponding to the transcript ID using biomart (38). Among the 16,548 differentially expressed transcripts identified, we were able to retrieve 8,386 genes identifiers.

### Gene ontology (GO) enrichment analysis

Go term enrichment of differentially expressed genes were determined using the web-based application Panther version 14.1 (39) according to the gene ID from Ensembl. We extracted GO slim terms defined according to the PANTHER classification system that were significantly over- and under-represented (according to statistical Fisher’s exact tests with FDR correction) in our comparisons in the three ontologies, Molecular Function, Cellular Component, and Biological Process.

### Estimation of the transposable element (TE) environment of each mouse gene

We downloaded the repeat-masker output file obtained on the latest version of the mouse genome assembly (http://hgdownload.cse.ucsc.edu/goldenpath/mm10/bigZips/). The file was parsed using the program one-code-to-find-them-all (40) with the --strict option to avoid false positive identification. In short, this tool allows to assemble each TE copy and provide their positions in the genome. For each annotated gene (54,067 genes), we determined all TE present inside the gene and in its 2-kb surrounding regions to cover the promoter region, and we computed the TE density and the TE coverage as proposed by Grégoire et al. (41). All genes were then clustered according to their level of TE density and TE coverage using the K-medoids algorithm as implemented in the pam() function of the R package (42). This method allows the unsupervised classification in a defined number of classes. After removing genes with no TE in their surrounding (corresponding to the gene category TE-free genes), the remaining genes were clustered using both density and coverage values to discriminate between the TE-very-poor (mean density of 8.167e-4 insertions/pb and mean coverage of 0.1296), TE-poor (mean density of 0.0019 insertions/pb and mean coverage of 0.3091), TE-rich (mean density of 0.0045 insertions/pb and mean coverage of 0.5747), and TE-very-rich genes (mean density of 0.1835 insertions/pb and mean coverage of 0.9877).

### Expression analysis of transposable elements (TEs)

To determine specifically the differentially expressed TE families between the two conditions, we used the TEtools pipeline (43) (https://github.com/l-modolo/TEtools). In this procedure, the reads were first aligned on all TE copies annotated in the mouse reference genome using Bowtie2 (44) with the most sensitive option and keeping a single alignment for reads mapping to multiple positions (–very-sensitive option). The TE sequences were retrieved using the program one-code-to-find-them-all (40). The read count was computed per TE family, adding all reads mapped on copies of the same family. The differential expression analysis of TE families between the two conditions was performed using DESeq2 (37) on the raw read counts, using the Benjamini–Hochberg multiple test correction (FDR level of 0.05 (45)).

### TE-initiated chimeric transcript detection

We determined the presence of TE-initiated chimeric transcripts in our dataset by using the pipeline LIONS (46). By providing the raw paired-end RNA-seq data for each condition, the genome sequence, the repeat masker output file and the gene annotation file (Mus_musculus.GRCm38.97.gtf), this tool is able to identify transcripts originated in a TE sequence in each sample and gives information concerning their positions relative to annotated genes.

### Statistical analyses

All statistical tests were performed using the R software version 3.6.2 (2019-12-12 (42)). Mosaic plots were generated using the mosaic.plot() function which allows the visualization of standardized residuals of a loglinear model considering a contingency table.

## Results

### Many differentially expressed genes between the two conditions

The Principal Component Analysis performed on normalized counts shows that normal and tumor samples are well discriminated on the second axis (31% of variation – Fig. 1A). We observed that 8,386 genes are differentially expressed between the normal and tumor conditions among the 54,067 genes annotated in the mouse genome. In more detail, 2,444 genes are down-regulated whereas 5,942 genes are up-regulated in the tumor condition when compared to the normal condition (Fig. 1B and Supplementary Table S1). We investigated among the differentially expressed genes, the proportion of genes according to their type (coding, non-coding, and pseudogenes). The large majority corresponds to coding genes (6,921 genes, 82.53%) whereas non-coding genes and as could be expected pseudogenes are much less represented (834 genes – 9.94% and 631 genes – 7.52%, respectively). On the contrary, these proportions are about the same in all categories when considering the genes that are not differentially expressed (14,208 coding genes – 31.10%, 14,421 non coding genes – 31.57%, and 17,052 pseudogenes – 37.33%).

**Fig. 1.**
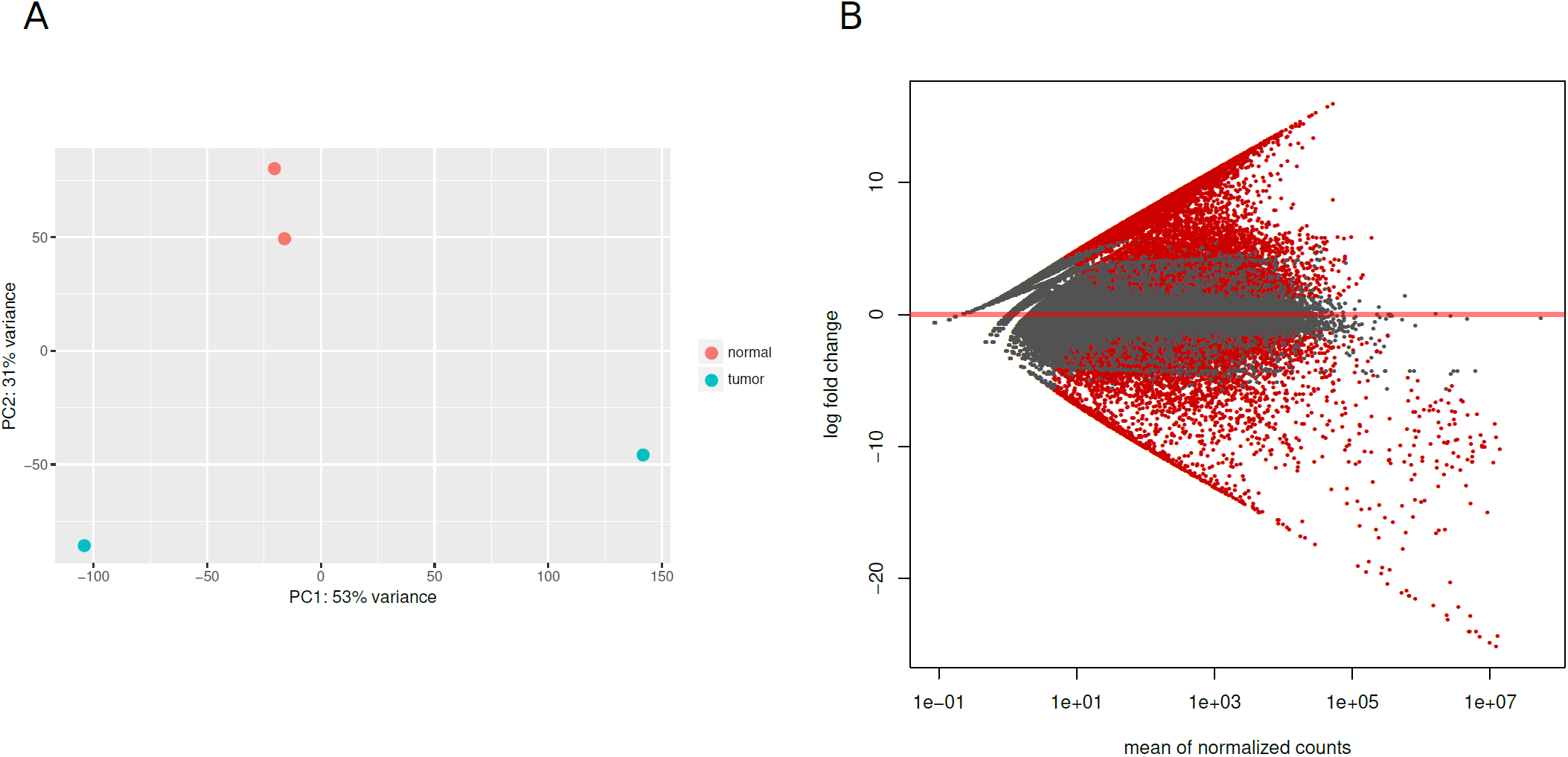
Differentially expressed genes between normal and tumor condition. (A) Principal Component (PC) Analysis performed for the samples using the gene expression values. The tumor and normal samples are well discriminated on the second axis. (B) MA plot representing the log2 fold change of expression between the two conditions on the y-axis and the mean of normalized read count the x-axis. Each dot represent a gene with the red dots representing differentially expressed genes at adjusted p-value < 0.1.

To determine whether the differentially expressed genes display particular functional enrichment, we performed a functional gene ontology (GO) analysis with the PANTHER V14.1 classification system. We compared the functions of the down- and the up-regulated genes to the complete gene set of the reference mouse genome. The differentially expressed genes show several profiles that differ according their up- or down-regulated status (Fig. 2, Supplementary Fig. S1 and Supplementary Table S2).

**Fig. 2.**
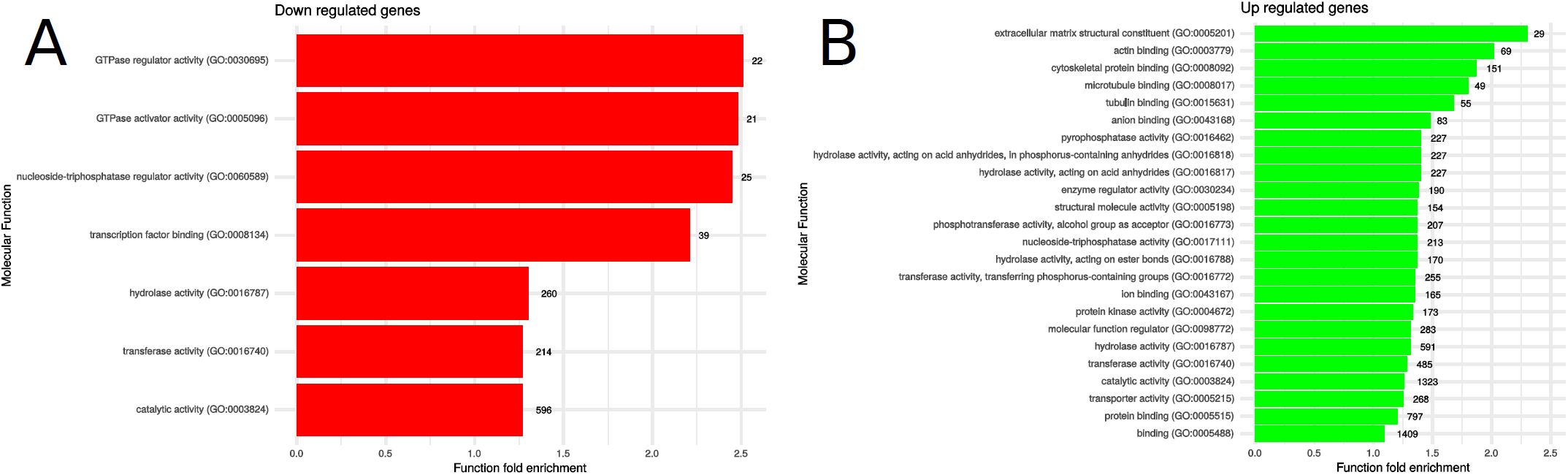
Molecular Function enrichment from GO analysis for down-(A) and up-regulated genes (B). Only function fold enrichment of more than 1 are represented.

Especially, the down-regulated genes are particularly enriched for GTPase (GO:0030695 and GO:0005096) and nucleoside-triphosphatase (GO:0060589) activities and in transcription factor binding (GO:0008134) since the number of down-regulated genes with these functions is twice as much as what is expected by chance (Fig. 2A). In the Biological Process, similarly, the down-regulated genes are particularly enriched in processes involved in endoplasmic reticulum (ER) to golgi vesicle-mediated transport (GO:0006888), organelle assembly (GO:0070925) and organonitrogen compound metabolic process (GO:1901564) (Supplementary Fig. S1A), which is congruent with the cell localization reflected by the Cellular Component GO. Indeed, the down-regulated genes are particularly enriched in genes whose function are localized near Golgi and ER compartments (Supplementary Fig. S1B). These results means that tumor pancreatic cells lack the products of these genes since they are down-regulated when compared to a normal cell state. When considering the up-regulated genes, we observed a functional enrichment different from those observed with the down-regulated genes, as could be expected. In the Molecular Function aspect for example, up-regulated genes are principally enriched in functions involved in extracellular matrix structural constituent (GO:0005201) and actin binding (GO:0003779) with twice as many genes as expected by chance, but also in the binding of cytoskeletal protein (GO:0008092), microtubule (GO0008017) and tubulin (GO:0015631) (Fig. 2B). The Biological Process classification indicates that up-regulated genes are enriched in the negative regulation of cell development (GO:0010721), the nuclear transport (GO:0051169), the organization of the extracellular matrix and structure (GO:0030198 and GO:0043062) the mitotic cell cycle phase transition (GO:0044772), the regulation of cell differentiation (GO:0045595) and the regulation of cell migration (GO:0030334) (Supplementary Fig. S1C). The main localization for the up-regulated genes as reflected by the Cellular Component classification indicated a bias toward extracellular matrix (GO:0031012) and cytoskeleton (GO:0015629 and GO:0005856) (Supplementary Fig. S1D).

### The TE environment near genes is associated with the gene expression divergence and with the gene type

We determined the TE environment, i.e. both the density and the coverage in TEs (see methods), of all mouse genes, including pseudogenes, coding and non-coding genes. All genes were clustered according to their TE environment (see methods) to identify different categories (Supplementary Table S1). We observed that the distributions corresponding to the number of genes according to their TE neighborhood and to the different types of genes (coding, non-coding, and pseudogenes) are different (Fig. 3, X-squared = 7134.3, df=8, p-value<2.2e-16). Indeed, coding genes display less than expected TE-free, TE-rich and TE-very-rich genes whereas TE-very-poor and TE-poor genes are over-represented. On the contrary, pseudogenes display more than expected TE-free, TE-rich and TE-very-rich genes. These results are consistent with what could be expected considering that natural selection will remove deleterious TEs near functional genes but not near non-functional ones. TEs are thus expected to accumulate more near pseudogenes. The non-coding genes display more than expected TE-free genes but lack TE-very-poor and TE-poor genes, which could be explained by their usual short length since they correspond to genes coding for small RNAs.

**Fig. 3.**
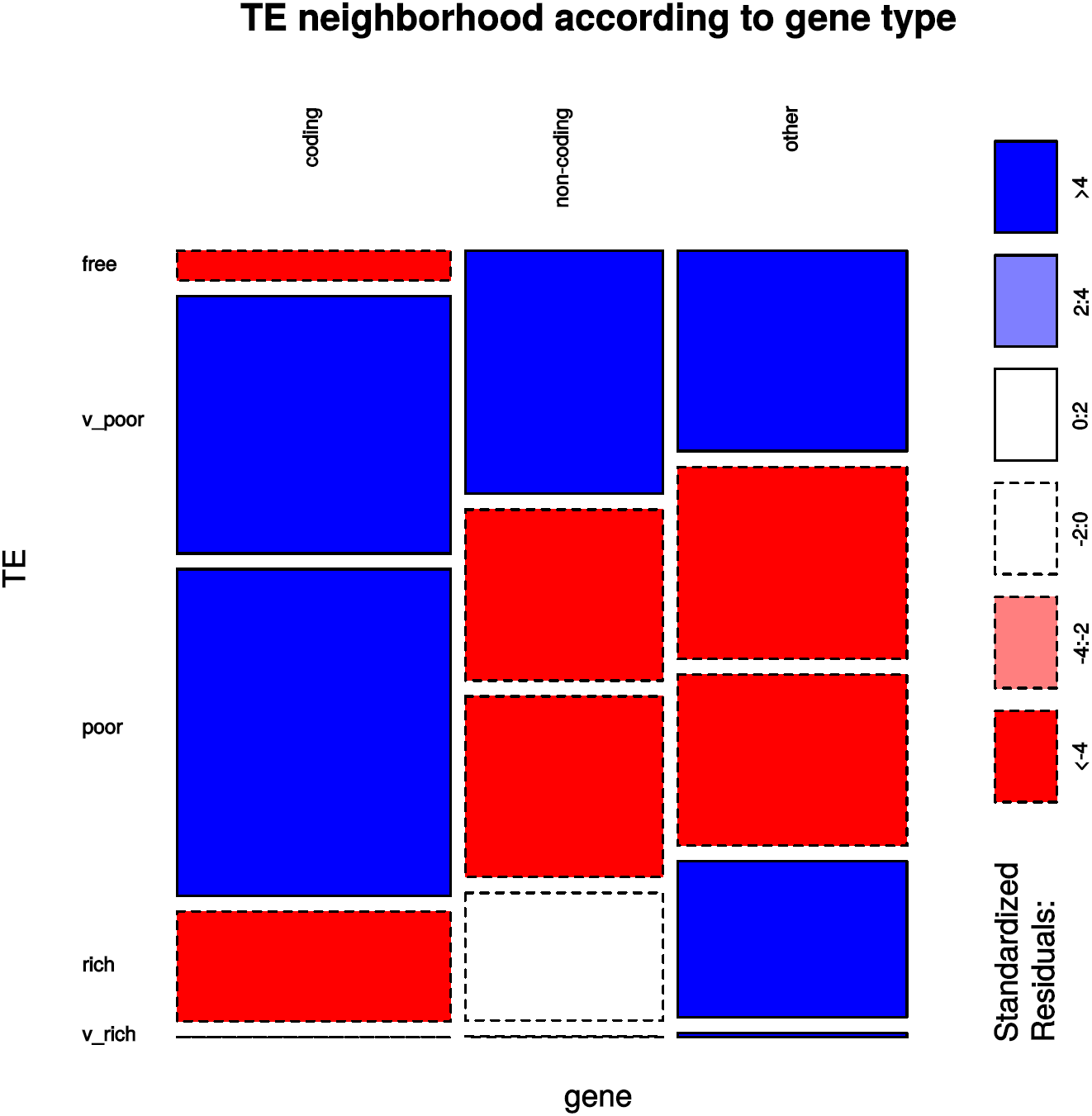
Mosaic-plot representing the proportion of genes according to their type (coding, non-coding or pseudogene) and to their TE environment (TE-free, TE-very-poor, TE-poor, TE-rich, and TE-very-rich). Blue boxes indicate that the observed proportions are more than what is expected whereas red boxes indicate that the observed proportions are less than what is expected.

We then wanted to determine whether the expression variation of genes between the two conditions is associated with their TE neighborhood. For that, we observed the distributions of differentially expressed genes according to their TE neighborhood and compared it to the distribution of genes with no expression divergence (Fig. 4).

**Fig. 4.**
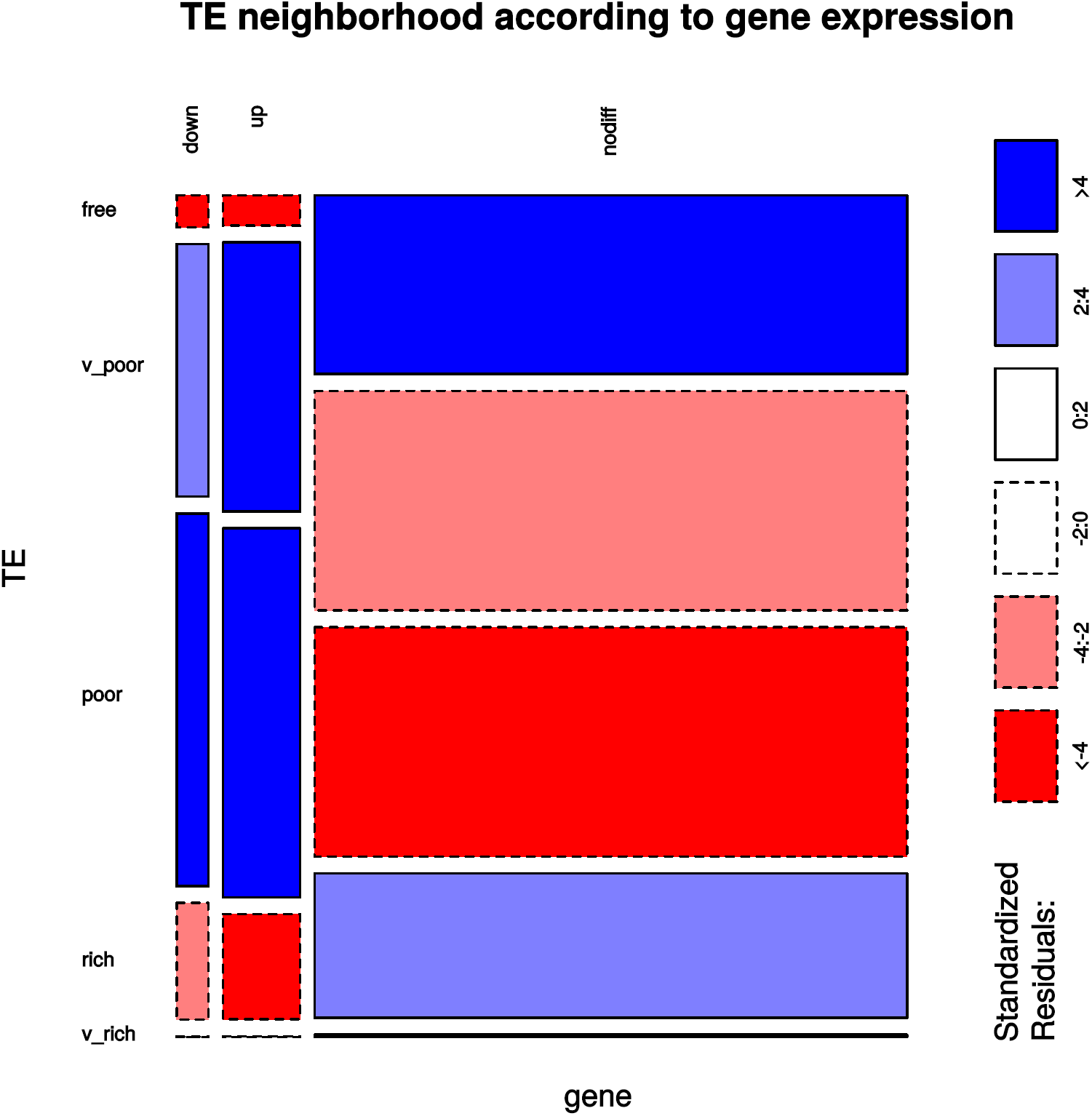
Mosaic-plot representing the proportion of genes according to their expression divergence between the two conditions (down-regulated, up-regulated or not-differentially expressed) and to their TE environment (TE-free, TE-very-poor, TE-poor, TE-rich, and TE-very-rich). Blue boxes indicate that the observed proportions are more than what is expected whereas red boxes indicate that the observed proportions are less than what is expected.

The results showed that the TE neighborhood is associated to the expression variation of the genes (X-squared = 2189.57, df=8, p-value<2.2e-16). Indeed, differentially expressed genes display an excess of TE-very-poor and TE-poor genes, whereas TE-free and TE-rich genes are under-represented. Genes with no differential expression show the opposite trend with an over-representation of TE-free genes and on a less extent of TE-rich genes. To determine if this trend is true whatever the type of genes, we did the same analyses separately for the pseudogenes, coding and non-coding genes. We first compared the differentially expressed coding genes versus coding genes with no expression variation to determine any association with the TE neighborhood. The distributions of the two set of genes are significantly different (X-squared = 311.2, df=4, p-value<2.2e-16) meaning that differentially expressed coding genes have different TE neighborhood when compared to coding genes with no expression divergence. In particular, there is an under-representation of TE-free and TE-rich genes and an over-representation of TE-poor genes among the differentially expressed coding genes (Supplementary Fig. S2). The non-coding genes and the pseudogenes also show different distributions when comparing differentially and not differentially expressed genes with an under-representation of TE-free genes and an over-representation of TE-poor genes in the differentially expressed genes (X-squared = 121.54, df=4, p-value<2.2e-16 for non-coding genes; X-squared = 54.63, df=4, p-value=3.88e-11 for pseudogenes; Supplementary Fig. S3 and S4). Thus, the TE neighborhood of genes differs according to the status of gene expression variation. Especially, TE-free genes are systematically under-represented among differentially expressed genes.

### Many families from all TE classes are differentially expressed

We looked more specifically at the TEs themselves to find out whether particular families present variations in their expression level between the two analyzed conditions. The Principal Component Analysis performed on normalized counts shows that normal and tumor samples are well discriminated on the first axis based on the TE expression (80% of variance) (Supplementary Fig. S5). This analysis allows us to show that among the 1,034 TE families annotated in the reference genome, 162 are significantly differentially expressed (Table 1 – Supplementary Fig. S6 to S9, Supplementary table S3). Among these families, 95 are down-regulated in tumor condition whereas 67 are up-regulated. We tested whether the number of families differentially expressed from each class (SINE, LINE, LTR and DNA) is representative of the whole genome. The results indicated a significant difference in the representation of each class with an excess of LTR retrotransposons and in a less extent of SINE elements whereas the DNA transposons are under-represented (Table 1; X-squared = 13.152, df=3, p-value=0.00432)

**Table 1:**
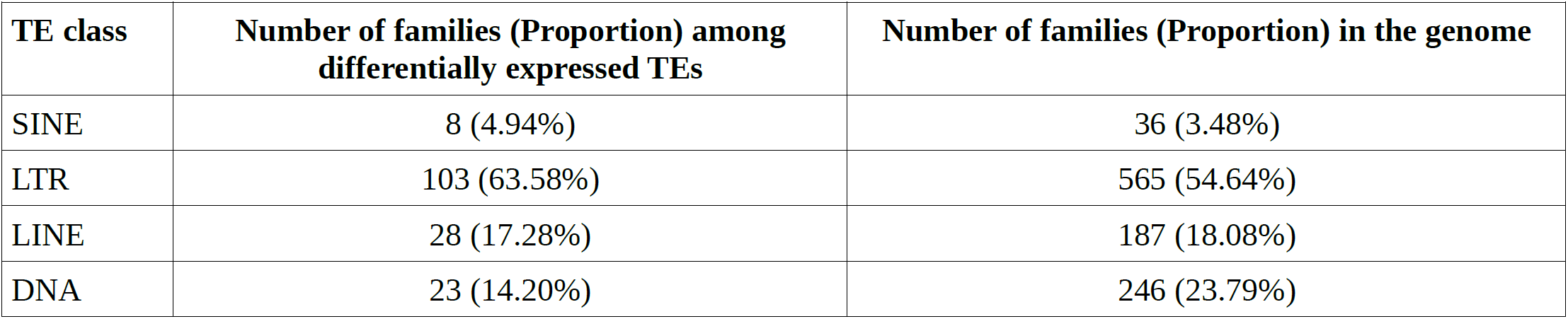
Number of TE families differentially expressed between normal and tumor condition compared to the all genome.

For the DNA transposons and the LINE elements, differentially expressed families belong to the most represented superfamilies in the genome (X-squared = 3.6336, df = 3, p-value = 0.3038 for the LINEs and X-squared = 2.217, df = 3, p-value = 0.5286 for the DNA transposons). In that respect, the majority of the differentially expressed LINE elements correspond to the L1 superfamily (Table 2; Supplementary Fig. S7). However, only three L1 families are up-regulated (Lx7, L1M3A, and L1MC5a) whereas 21 are down-regulated. Among the three L2 families to be differentially expressed, only L2d is up-regulated. In the DNA transposon class, elements from only two superfamilies (over a total in the genome of seven) are differentially expressed. Among them, elements from the hAT superfamily are the most numerous compared to the TcMar superfamily (Table 2; Supplementary Fig. S8). However, there are comparatively more TcMar families to be up-regulated in the cancer condition. In the case of SINE elements and LTR retrotranposons, the picture is not the same (X-squared = 14.859, df = 5, p-value = 0.01098 for the LTR retrotransposons and X-squared = 10.943, df = 3, p-value = 0.01204 for the SINEs). This indicates that the number of differentially expressed families is not dependent on the proportion of their corresponding superfamily in the genome. More specifically, the B2 and B4 superfamilies are particularly over-represented among the differentially expressed families (standardized residual = 3.26). For the LTR-retrotransposons, ERVK superfamilies are particularly over-represented (standardized residual = 3.08) whereas ERVL are very under-represented among the differentially expressed families (standardized residual = −2.1). In more details, the majority of SINE families from three superfamilies are down-regulated (B2, B4 and tRNA) whereas only one is up-regulated (in the Alu superfamily) (Table 2; Supplementary Fig. S9). Among the LTR retrotransposons, different families of the same superfamilies are either down- or up-regulated (ERV1, ERVK, ERVL, and ERVL-MalR; Table 2; Supplementary Fig. S6). The superfamilies ERV1 and ERVK include the largest amount of differentially expressed individual families. Interestingly, the majority of ERV1 elements are up-regulated in tumor condition, whereas there is about the same proportion of families down- or up-regulated belonging to the ERVK superfamily. Globally, as can be expected, the youngest families, which are also those that are likely to be still active, correspond to the families presenting a differential expression.

**Table 2 :** Number of TE families according to their class and superfamily.

| Class | Superfamilies | In the genome | Down-regulated | Up-regulated |
| --- | --- | --- | --- | --- |
| DNA | hAT | 139 | 8 | 6 |
|  | TcMar | 70 | 3 | 5 |
|  | Kolobock | 1 | 0 | 0 |
|  | PIF Harbinger | 1 | 0 | 0 |
|  | PiggyBac | 2 | 0 | 0 |
|  | RC | 3 | 0 | 0 |
|  | Unknown | 30 | 1 | 0 |
| LINE | CR1 | 19 | 1 | 0 |
|  | DongR4 | 1 | 0 | 0 |
|  | Jockey | 1 | 0 | 0 |
|  | L1 | 149 | 21 | 3 |
|  | L2 | 11 | 2 | 1 |
|  | Penelope | 1 | 0 | 0 |
|  | RTE | 5 | 0 | 0 |
| LTR | ERV1 | 107 | 4 | 14 |
|  | **ERVK\*** | 260 | 35 | 28 |
|  | ERVL<sup>§</sup> | 106 | 6 | 5 |
|  | ERVL-MaLR | 59 | 7 | 4 |
|  | Gypsy<sup>§</sup> | 20 | 0 | 0 |
|  | Unknown<sup>§</sup> | 13 | 0 | 0 |
| SINE | 5SDeuL2 | 1 | 0 | 0 |
|  | Alu<sup>§</sup> | 15 | 0 | 1 |
|  | **B2\*** | 5 | 3 | 0 |
|  | **B4\*** | 4 | 3 | 0 |
|  | ID | 3 | 0 | 0 |
| | MIR\$ | 5 | 0 | 0 |
|  | tRNA | 3 | 1 | 0 |

\*over-represented among the differentially expressed TE families
§under-represented among the differentially expressed TE families

### Few TE initiated chimeric transcripts in cancer condition but involving differentially expressed genes

Since TEs are known to provide alternative promoters to genes, especially in cancer condition, we have investigated the presence of chimeric gene transcripts originated in TE sequences. Using the complete RNAseq dataset, we identified 189 TE-initiated transcripts. Among these 189 chimeric transcripts, 24 are found exclusively in the two tumor datasets, whereas 156 are found in either one of the tumor datasets but not in the two normal datasets. Among the TEs involved in chimeric transcripts, the majority corresponds to LTR retrotransposons and LINEs (18 elements over 24) (Table 3).

**Table 3 :** TE-initiated transcripts present in tumor datasets exclusively.

| Transcript ID | Exon number | Ref ID | Ensembl ID | Strand relationship* | Repeat Name | Position |
| --- | --- | --- | --- | --- | --- | --- |
| Sou2_1.1579:Sou2_1.1579.1 | 2 | Chit1 | ENSMUSG00000026450 | s | ORR1A4:LTR:ERVL-MaLR | chr1:134111126-134138772 |
| Sou2_1.1755:Sou2_1.1755.3 | 10 | Mir7682; Trmt1l | ENSMUSG00000053286 | s | L1_Mus1:LINE:L1 | chr1:151451665-151452238 |
| Sou2_1.2094:Sou2_1.2094.1 | 1 | Uap1 | ENSMUSG00000026670 | as | RMER5:LTR:ERV1 | chr1:170139905-170141570 |
| Sou2_1.2553:Sou2_1.2553.1 | 1 | . |  | i | Lx10:LINE:L1 | chr1:191817136-191817755 |
| Sou2_1.4269:Sou2_1.4269.1 | 1 | Mgat4c | ENSMUSG00000019888 | s | RLTR1A2_MM:LTR:ERV1 | chr10:102374053-102374644 |
| Sou2_1.2976:Sou2_1.2976.1 | 1 | LOC108167355; H60b | ENSMUSG00000075297 | u | L1MA4:LINE:L1 | chr10:22272619-22463112 |
| Sou2_1.11563:Sou2_1.11563.1 | 1 | . |  | i | RLTR45:LTR:ERVK | chr14:13749178-13749726 |
| Sou2_1.17592:Sou2_1.17592.1 | 1 | H2-Q1 | ENSMUSG00000079507 | u | RLTR13D3A1:LTR:ERVK | chr17:35323491-35346865 |
| Sou2_1.18014:Sou2_1.18014.1 | 7 | Pja2 | ENSMUSG00000024083 | s | Charlie7:DNA:hAT-Charlie | chr17:64308622-64313188 |
| Sou2_1.20062:Sou2_1.20062.1 | 9 | Lgals12 | ENSMUSG00000024972 | s | MT2A:LTR:ERVL | chr19:7606662-7610512 |
| Sou2_1.22396:Sou2_1.22396.2 | 2 | LOC108168773 |  | as | RLTR47:LTR:ERVK | chr2:78347393-78413343 |
| Sou2_1.26436:Sou2_1.26436.1 | 14 | Fam73a | ENSMUSG00000054942 | s | B1_Mur1:SINE:Alu | chr3:152335252-152338059 |
| Sou2_1.27831:Sou2_1.27831.1 | 2 | Mmachc | ENSMUSG00000028690 | s | B1_Mus2:SINE:Alu | chr4:116704507-116706179 |
| Sou2_1.26951:Sou2_1.26951.1 | 1 | Galt; Sigmar1; Il11ra1 | ENSMUSG00000036073 ;ENSMUSG00000036078 ;ENSMUSG00000073889 | c | B2_Mm2:SINE:B2 | chr4:41754014-41755419 |
| Sou2_1.27439:Sou2_1.27439.1 | 3 | Gm12602 | ENSMUSG00000085569 | s | B4A:SINE:B4 | chr4:89127183-89127467 |
| Sou2_1.30046:Sou2_1.30046.1 | 3 | Stim2 | ENSMUSG00000039156 | s | UCON29:DNA:PiggyBac? | chr5:54053371-54075394 |
| Sou2_1.32628:Sou2_1.32628.1 | 1 | . |  | u | MMVL30-int:LTR:ERV1 | chr6:41196263-148638534 |
| Sou2_1.36054:Sou2_1.36054.1 | 2 | Vmn2r79 | ENSMUSG00000090362 | s | MuRRS4-int:LTR:ERV1 | chr7:87023002-87038161 |
| Sou2_1.39719:Sou2_1.39719.1 | 1 | Ntpcr | ENSMUSG00000031851 | s | MTEa:LTR:ERVL-MaLR | chr8:125729767-125730147 |
| Sou2_1.37729:Sou2_1.37729.2 | 1 | Mrps31 | ENSMUSG00000031533 | s | RMER5:LTR:ERV1 | chr8:22410432-22411584 |
| Sou2_1.42166:Sou2_1.42166.1 | 1 | Gm39469;<br>2010315B03Rik | ENSMUSG00000110834 ;<br>ENSMUSG00000074829 | s | L1Md_F2:LINE:L1 | chr9:124153298-124328369 |
| Sou2_1.43348:Sou2_1.43348.2 | 2 | Morf4l2 | ENSMUSG00000031422 | s | L1MC5:LINE:L1 | chrX:136739805-136740236 |
| Sou2_1.42242:Sou2_1.42242.1 | 1 | 2010204K13Rik | ENSMUSG00000063018 | s | RMER4B:LTR:ERVK | chrX:7412153-7412661 |
| Sou2_1.42316:Sou2_1.42316.1 | 1 | Ebp | ENSMUSG00000031168 | s | L1MEd:LINE:L1 | chrX:8185401-8190553 |

\*as: anti-sense long non-coding RNA; s: sense; c: complex (involving multiple genes); u: undetermined; i: intergenic

We thus looked into the 24 chimeric transcripts expressed in the two tumor datasets in more details and we were able to determine that four of them are associated with no gene, four are associated with more than one gene and 16 are associated to a single gene. To find out to which expression category these genes belong, we retrieved the ENSEMBL identifiers, which led to 23 genes. Among them, 11 are up-regulated in tumor condition, whereas three are down-regulated, the remaining nine do not show any expression divergence. This distribution is statistically different from what is observed when considering all genes (5,942 up-regulated genes, 2,444 down-regulated genes, and 45,681 genes with no differential expression on the total dataset; X-squared = 95.727, df=2, p-value<2.2e-16) meaning that there is an excess of differentially expressed genes among those presenting a transcription initiation inside a TE sequence. For example, the gene ENSMUSG00000031422 (mortality factor 4 like 2 (Morf4l2)), which is up-regulated in the tumor condition, contains a LINE element in its last intron which is responsible for the transcription of a chimeric sequence. Gene functional classifications were determined using the PANTHER V14.1 software and allowed us to determine that the majority of these genes are involved in binding (GO:0005488) and catalytic activity (GO:0003824), mainly located inside the cell (GO:0005623) and in processes corresponding to biological regulation (GO:0065007), metabolic process (GO:0008152), and cellular process (GO:0002376) (Supplementary Fig. S10).

## Discussion

In this work, we have investigated the expression divergence of genes between tumor and normal conditions of pancreatic tissues from mice to determine whether the presence of TEs in the neighborhood of genes may be associated with any expression variation.

We observed that about 15% of the mouse genes are deregulated between the normal and tumor conditions. The present analysis was performed with only two replicates per sample, which allows statistical assessment of differential gene expression. It has been proposed that having more than 20 replicates would allow the detection of more than 85% of all differentially expressed genes independently of their fold change (47). However, this may not be biologically relevant for very low values. In our analyses we used a very stringent p-value threshold along with one of the best tool to perform such analyses with a small number of replicates (47) which guaranties a good confidence in the differentially genes identified. Another confounding effect could be the difference of genetic background between mice for each condition. Indeed, it has been shown that substantial sequence differences exist between balb/C and the reference mouse genome, in particular, more than 30,000 unique SNPs have been identified (48). When considering all laboratory strains compared to the reference genome, it has been shown that allele-specific variation could be responsible of tissue-specific expression bias for 12% of transcripts (48). Then some of the observed variation could be due to this effect rather than real differences between conditions. We could thus expect to observe no functional bias in this case, assuming that all genes can be impacted. Nevertheless, up- and down-regulated genes showed involvement in different functions as demonstrated by GO analyses. Some of these functions were already mentioned in transcriptional analyses performed on pancreatic human samples but only in the case of up-regulated genes, which can be explained by the small sample size in the human analysis (49). However, when comparing individual genes and their expression status, we were able to identify the same up- and down-regulated genes. We also identified similar up- and down-regulated genes as those identified in another work performed also on human pancreatic data but focusing on known genes to be involved in pancreatic tumor progression (50). In addition to coding genes, we identified as well non-coding genes that were differentially expressed between the two conditions. These genes are usually coding for regulatory non-coding RNAs like micro-RNAs (miRNA) or long-non-coding RNAs (lncRNA). Such genes have already been described as having a role in the development and progression of pancreatic cancer (51,52). Although we identified only one miRNA (miR155) known to be up-regulated in the human pancreatic cancer among our dataset, we also identified in the differentially expressed non-coding genes, three lncRNAs, Morrbid, Malat1 (both up-regulated) and Trp53cor1 (down-regulated), known to be implicated in several other cancers in human (53–55). These results show that the mouse pancreatic model is indeed very representative of the human disease in term of gene expression.

We then wanted to determine whether there may be an association between the expression divergence of genes between the two conditions and their neighborhood in TEs. For that, we classified each gene according to the proportion of TEs that can be identified in and around them. Our results showed that TE-poor and TE-very-poor-genes are over-represented among differentially expressed genes whereas TE-free genes are largely under-represented, implying that the presence of TEs near genes may be associated to their expression divergence. The presence of TEs has already been suspected to influence the expression of genes near which they are inserted. In particular, TEs are considered as a genome-wide source of regulatory elements (see for a review Chuong et al. (56)). In addition, new insertions of LTR-retrotransposons and SINEs have been shown to affect gene expression variation between mouse and rat, indicating a major impact on the evolution of gene expression levels in rodents (57). More recently, TEs have been shown to provide CTCF binding site divergence between human and mouse that explains a large fraction of variable chromatin loops corresponding to gene expression variability (58). The effect of TEs on gene expression is particularly exemplified when comparing cancer and normal condition. Indeed, in previous studies in human we showed that the expression level of TE-free genes was lower that TE-rich genes, particularly in tumor tissues (29,59). Recently, the characterization of the global profile of TE onco-exaptation, which is the reactivation of TEs influencing the oncogenesis, highlighted this phenomenon as an important mechanism for neighboring oncogene activation (30). This can be explained by the fact that TEs are usually deregulated in cancer condition, especially via the loss of DNA methylation or of repressing histone modifications (see for a review Anwar et al. (60). Hence, epigenetic modifications occurring on TEs may influence neighboring genes. For example, we already demonstrated in human that genes with TEs in their neighborhood are more likely to observe a change in their associated epigenetic modifications when comparing normal and tumor condition than TE-free genes (41).

We were able to identify 162 TE families that are deregulated in cancer condition in the mouse, of which about 40% are up-regulated. Among these deregulated TE families, we found a high proportion of retrotransposons from the superfamilies L1, ERV1, ERVL/MaLR and especially ERVK. This is congruent with what we know concerning the active TEs in the mouse genome. Indeed, several active superfamilies from the three described classes of endogenous retroviruses present in the mouse genome have been described (IAP, MusD/ETn, MMTV, MuERV-L, MaLR, MuLV, MuRRS, GLN, and VL30) of which five correspond to up-regulated TEs in our analyzed cancer condition (61). Interestingly, the IAP and ETn elements are known to be responsible for the majority of new insertions observed in mouse germ line (62). Among the up-regulated LTR retrotransposons, we also identified the RLTR13D6 element from the ERVK superfamily. This element has been shown to harbor multiple transcription binding sites, indicating that this element is susceptible to influence the expression of neighboring genes (63). The number of active L1 non-LTR retrotransposons has been estimated to be largely higher than in human, with three known active subfamilies (64). Globally, we were able to identify several up-regulated TE families that may contribute to the cancer progression by various factors that could be mutational insertion or gene expression variation.

We detected a small number of chimeric TE-initiated transcripts in our dataset among which a majority corresponds to differentially expressed genes. This type of chimeric transcripts originates when the transcription starts at a TE promoter and encompass a neighboring gene. This phenomenon may result in the production of a new chimeric ORF which may acquire a new oncogenic potential (see for a review Babaian and Mager (18)). For example, a study performed on diffuse large B-cell lymphoma data in human leads to the identification of 98 TE-chimeric gene transcripts (65). Among them, one was shown to encode a novel chimeric isoform with new characteristics that may contribute to the pathogenesis of this cancer. This illustrates the most direct effect that TEs may have on genes that would deserve more investigation in cancer research.

To conclude, in this study, we have shown that the presence of TEs near genes is associated to the gene differential expression in the tumor condition. These results highlight again the potential role of TEs if not in the cancer initiation, at least in the global cellular disturbance occurring in cancer cells. However, a limitation is that it was not possible to evaluate the individual role of new TE insertions occurring in the tumor genomes with our data. Indeed, new TE insertions could also be responsible of important gene expression variation. Future investigations thus need to be done in that direction.

## Acknowledgments

We thank the LBBE for the financial support and L. Bartholin for the gift of pancreatic tumor tissues from mice. This work was performed using the computing facilities of the CC LBBE/PRABI.

## Author contributions

EL conceived the study, performed the identification of TE neighborhood of gene and the differential expression analyses of TEs and genes, and wrote the first draft of the manuscript; NB performed RNA extraction and library preparation; VN conducted the chimeric transcript analyses; CN contributed to the collegial construction of the standards of science, by developing the methodological framework, the state-of-the-art, and by ensuring post-publication follow-up; all authors reviewed and edited the different versions of the manuscript.

## Supporting Information

**Supplementary Fig. S1:** Function enrichment from GO analysis for down- and up-regulated genes. Only function fold enrichment of more than 1 are represented. (A) Biological Process enrichment for down-regulated genes. (B) Cellular Component enrichment for down-regulated genes (C) Biological Process enrichment for up-regulated genes. (D) Cellular Component enrichment for up-regulated genes.

**Supplementary Fig. S2**: Mosaic-plot representing the proportion of coding genes according to their expression divergence (up-regulated or not-differentially expressed) and to their TE environment (TE-free, TE-very-poor, TE-poor, TE-rich, and TE-very-rich). Blue boxes indicate that the observed proportions are more than what is expected whereas red boxes indicated that the observed proportions are less than what is expected.

**Supplementary Fig. S3**: Mosaic-plot representing the proportion of non-coding genes according to their expression divergence (up-regulated or not-differentially expressed) and to their TE environment (TE-free, TE-very-poor, TE-poor, TE-rich, and TE-very-rich). Blue boxes indicate that the observed proportions are more than what is expected whereas red boxes indicated that the observed proportions are less than whas is expected.

**Supplementary Fig. S4**: Mosaic-plot representing the proportion of pseudogenes according to their expression divergence (up-regulated or not-differentially expressed) and to their TE environment (TE-free, TE-very-poor, TE-poor, TE-rich, and TE-very-rich). Blue boxes indicate that the observed proportions are more than what is expected whereas red boxes indicated that the observed proportions are less than what is expected.

**Supplementary Fig. S5**: Principal Component Analysis performed on normalized counts of TE expression. Data from tumor condition are in red whereas data from normal condition are in blue. Circles correspond to the first replicate of each data and Triangles correspond to the second replicates. The two conditions are well discriminated on the first axis of the PCA.

**Supplementary Fig. S6**: Log2fold change of differentially expressed families from the LTR retrotransposon class according to their superfamily. Red and blue bars indicate respectively down- and up-regulation.

**Supplementary Fig. S7**: Log2fold change of differentially expressed families from the LINE retrotransposon class according to their superfamily. Red and blue bars indicate respectively down- and up-regulation.

**Supplementary Fig. S8**: Log2fold change of differentially expressed families from the DNA transposon class Red and blue bars indicate respectively down- and up-regulation.

**Supplementary Fig. S9**: Log2fold change of differentially expressed families from the SINE non-LTR retrotransposon class. Red and blue bars indicate respectively down- and up-regulation.

**Supplementary Fig. S10**: Function enrichment from GO analysis for chimeric transcript initiated in TEs that are differentially expressed between tumor and normal condition

**Supplementary table S1:** Differentially expressed genes identified between the normal and tumor condition associated to their TE density and coverage (globally and for each TE class)

**Supplementary table S2:** GO analyses for the differentially expressed genes

**Supplementary table S3:** Differentially expressed TEs according to their families.

## Notes

### Competing Interest Statement

The authors have declared no competing interest.

