## Supplementary material for "Transposable element activity in the transcriptomic analysis of mouse pancreatic tumors": Supplementary_figures.pdf

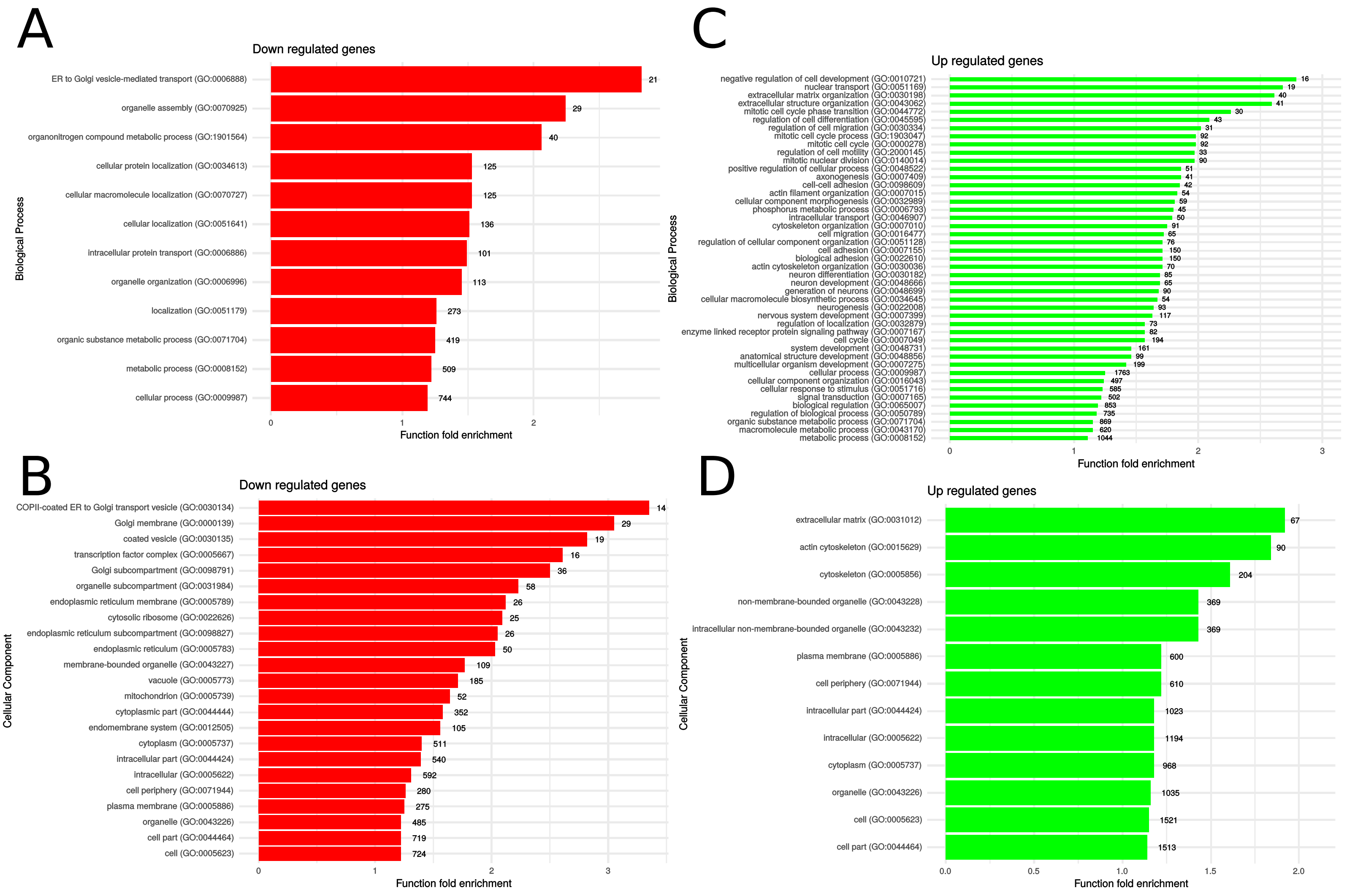


**Supplementary Fig. S1**: Function enrichment from GO analysis for down- and up-regulated genes. Only function fold enrichment of more than 1 are represented. (A) Biological Process enrichment for down-regulated genes. (B) Cellular Component enrichment for down-regulated genes (C) Biological Process enrichment for up-regulated genes. (D) Cellular Component enrichment for up-regulated genes.

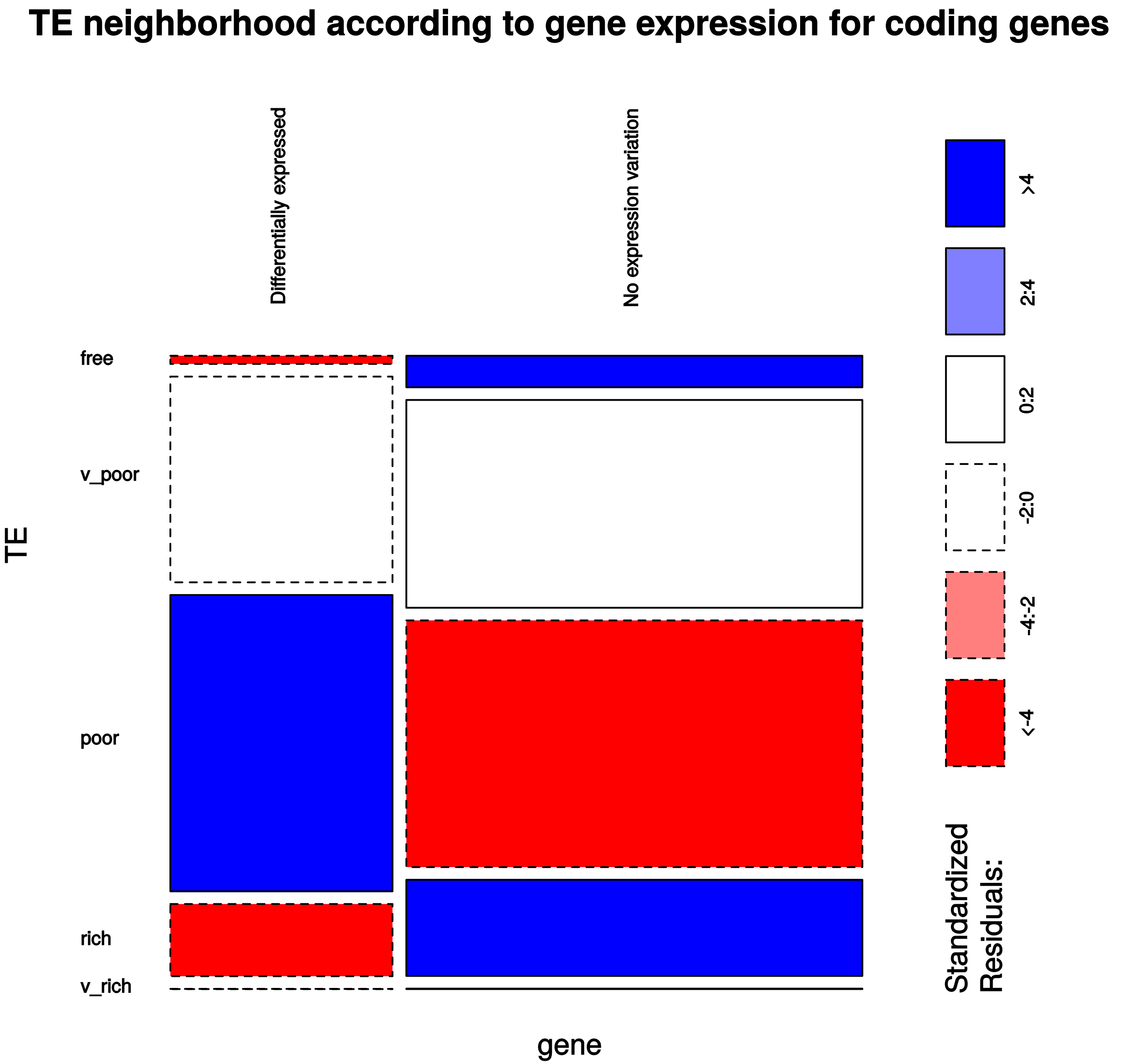


**Supplementary Fig. S2**: Mosaic-plot representing the proportion of coding genes according to their expression divergence (up-regulated or not-differentially expressed) and to their TE environment (TE-free, TE-very-poor, TE-poor, TE-rich, and TE-very-rich). Blue boxes indicate that the observed proportions are more than what is expected whereas red boxes indicated that the observed proportions are less than whas is expected.

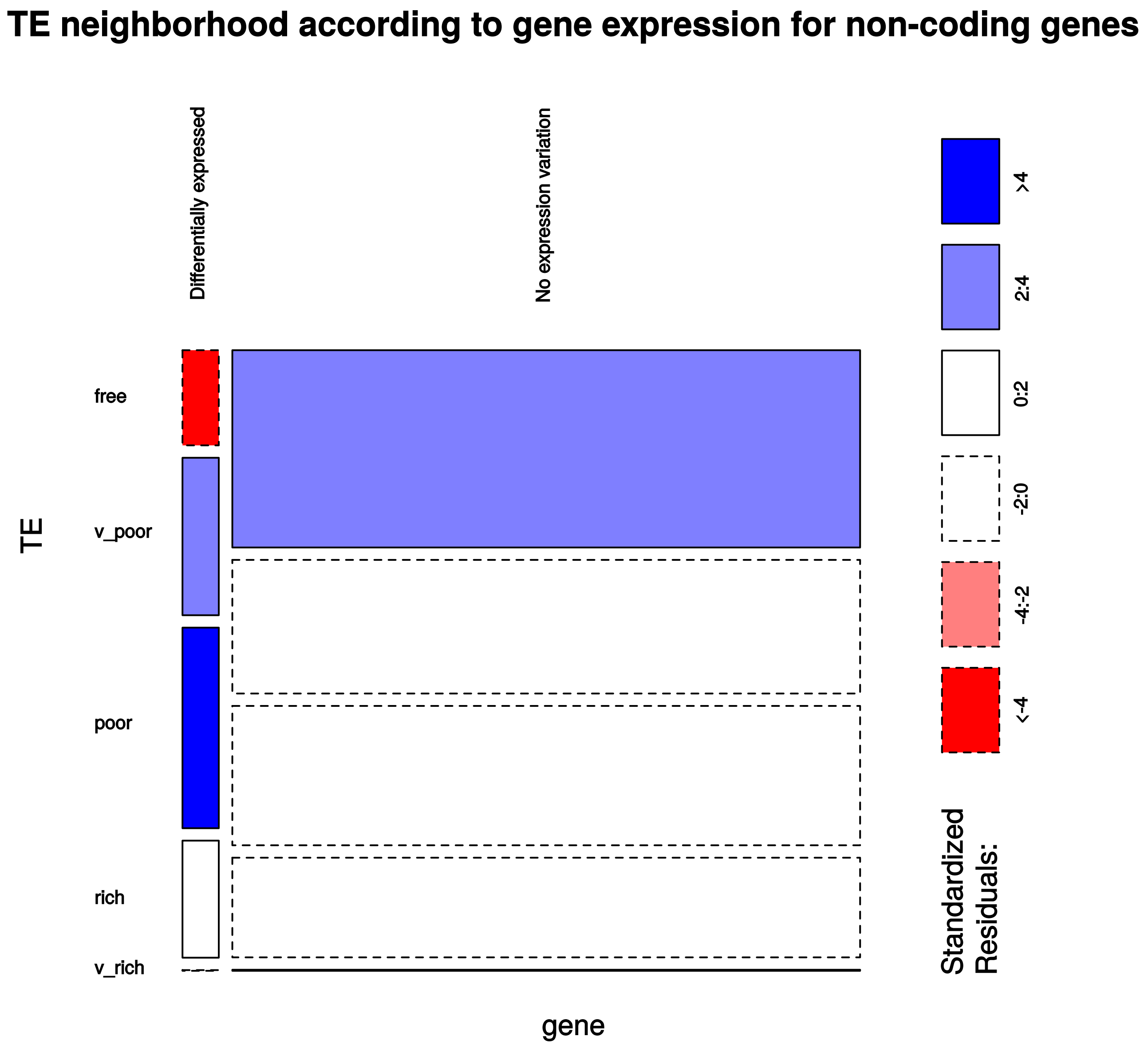


**Supplementary Fig. S3**: Mosaic-plot representing the proportion of non-coding genes according to their expression divergence (up-regulated or not-differentially expressed) and to their TE environment (TE-free, TE-very-poor, TE-poor, TE-rich, and TE-very-rich). Blue boxes indicate that the observed proportions are more than what is expected whereas red boxes indicated that the observed proportions are less than whas is expected.

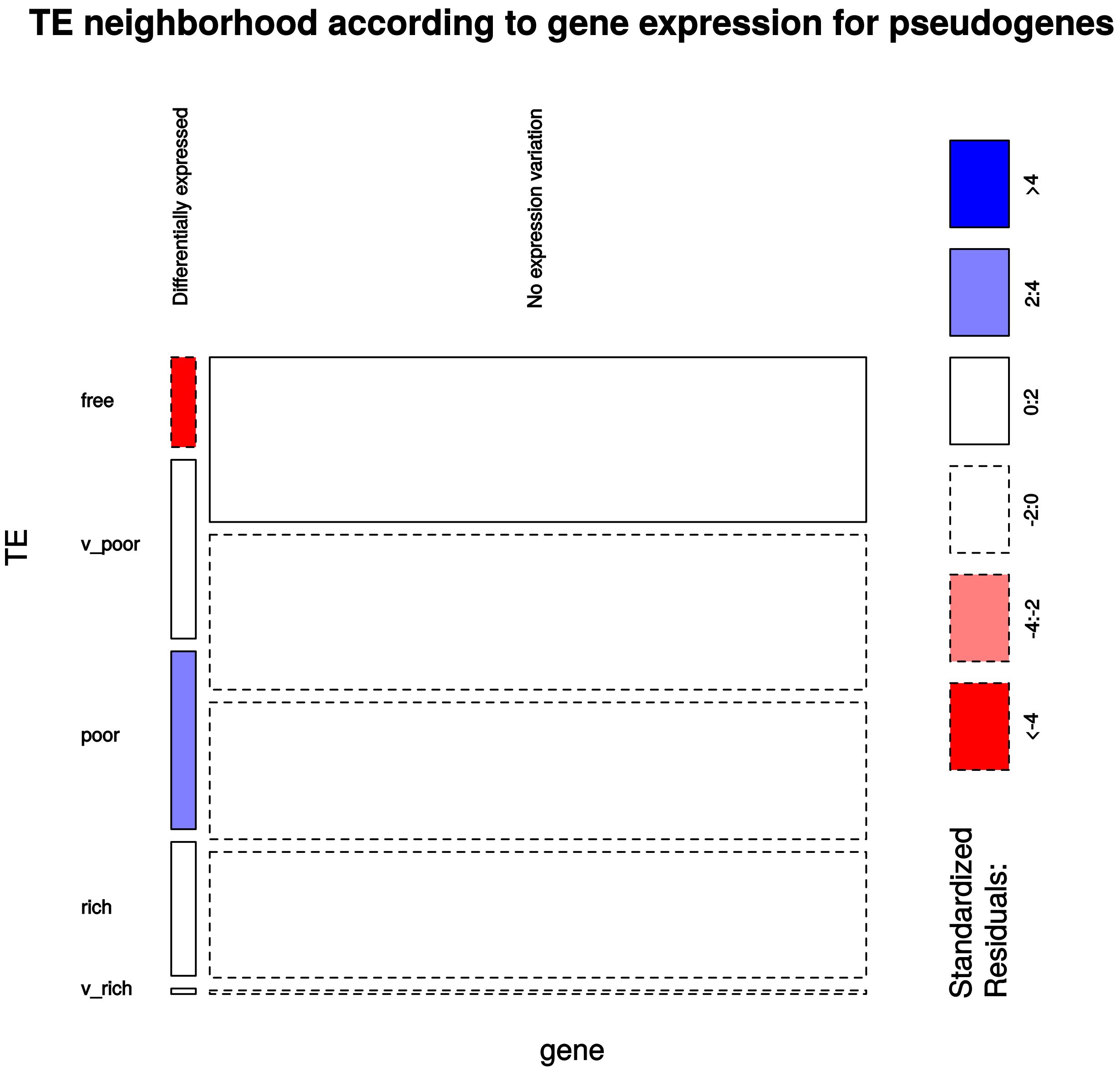


**Supplementary Fig. S4**: Mosaic-plot representing the proportion of pseudogenes according to their expression divergence (up-regulated or not-differentially expressed) and to their TE environment (TE-free, TE-very-poor, TE-poor, TE-rich, and TE-very-rich). Blue boxes indicate that the observed proportions are more than what is expected whereas red boxes indicated that the observed proportions are less than whas is expected.

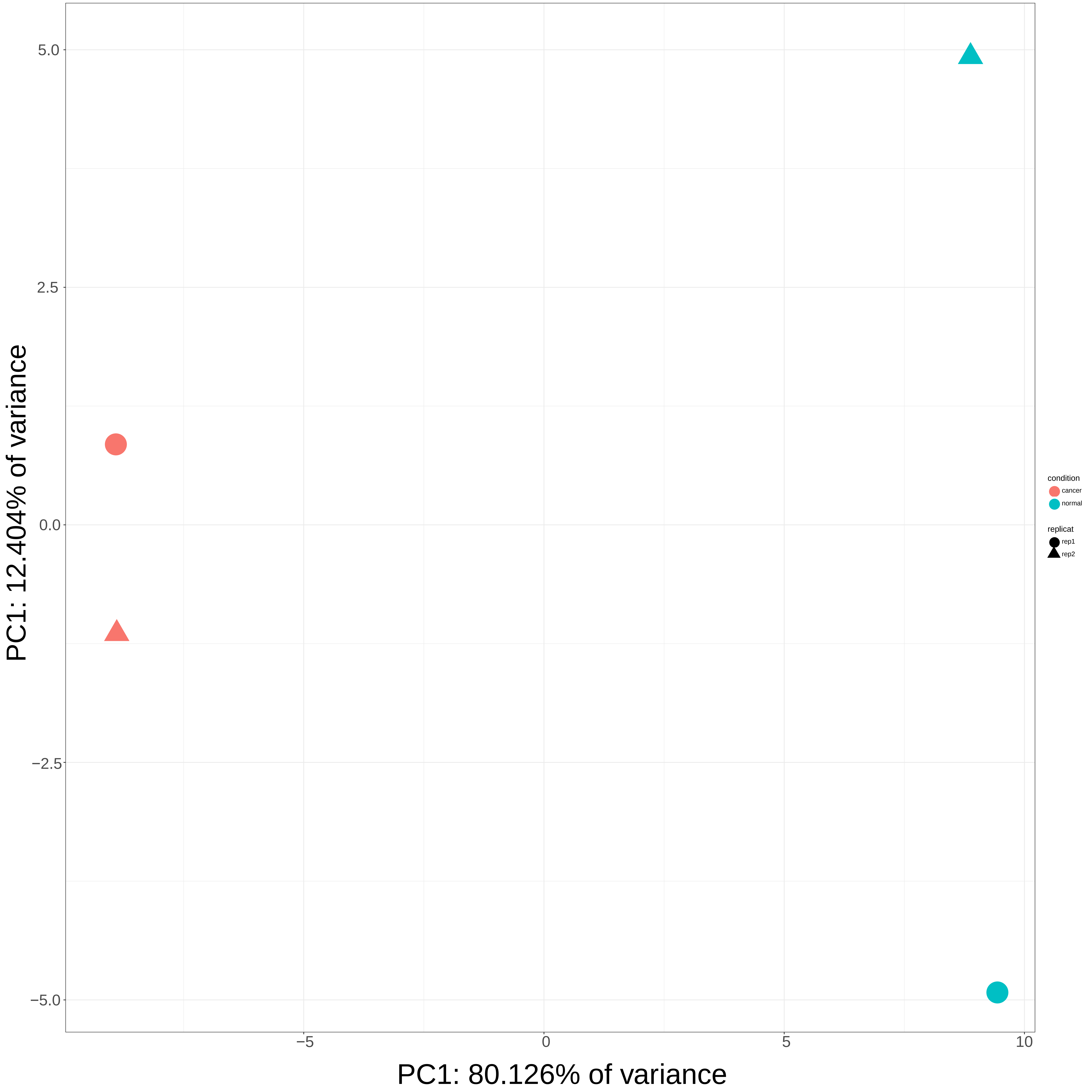


**Supplementary Fig. S5**: Principal Component Analysis performed on normalized counts of TE expression. Data from tumor condition are in red whereas data from normal condition are in blue. Circles correspond to the first replicate of each data and Trianges correspond to the second replicates. The two conditions are well discriminated on the first axis of the PCA.

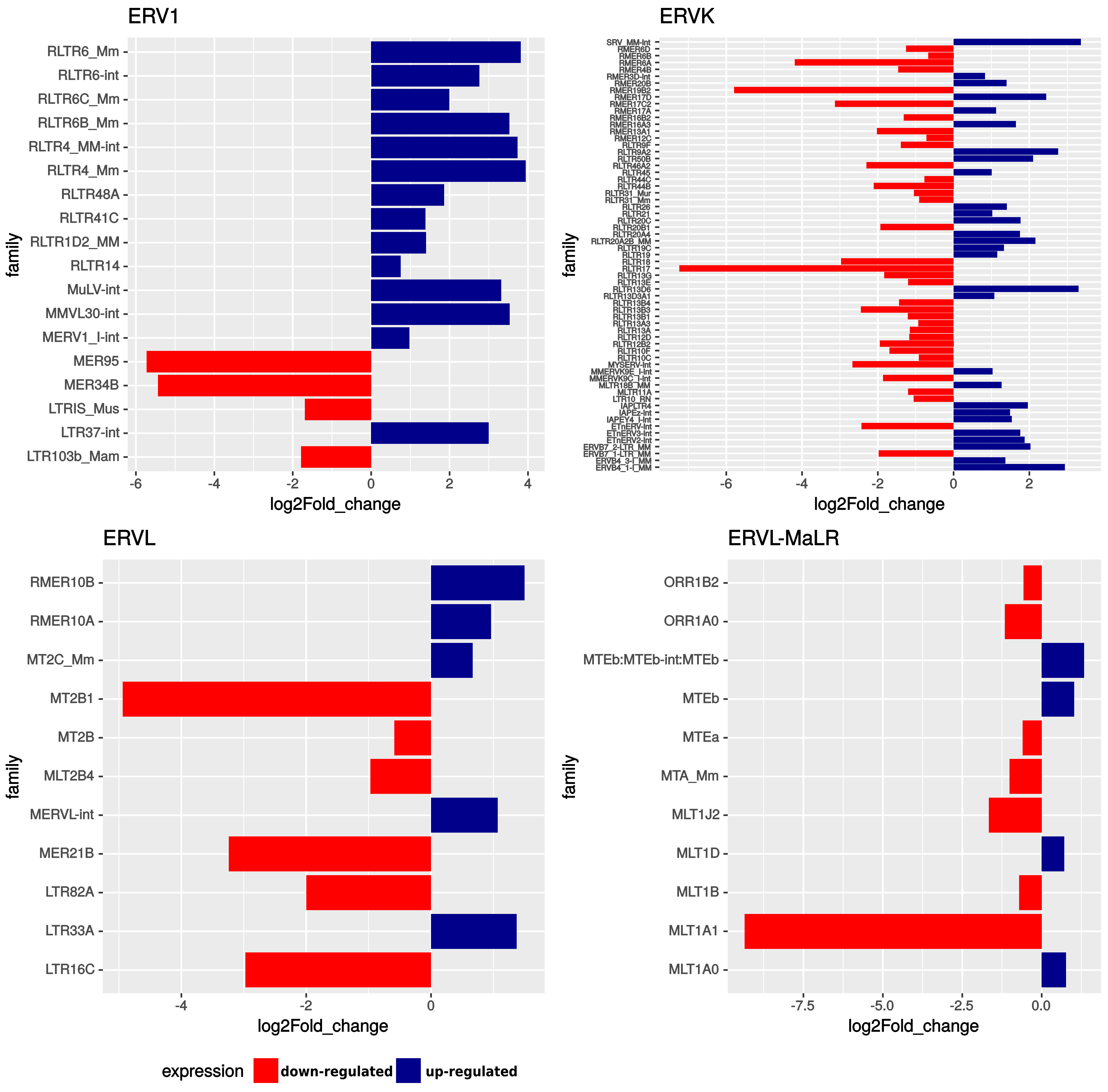


**Supplementary Fig. S6:** Log2fold change of differentially expressed families from the LTR retrotransposon class according to their superfamily. Red and blue bars indicate respectively down- and up-regulation.

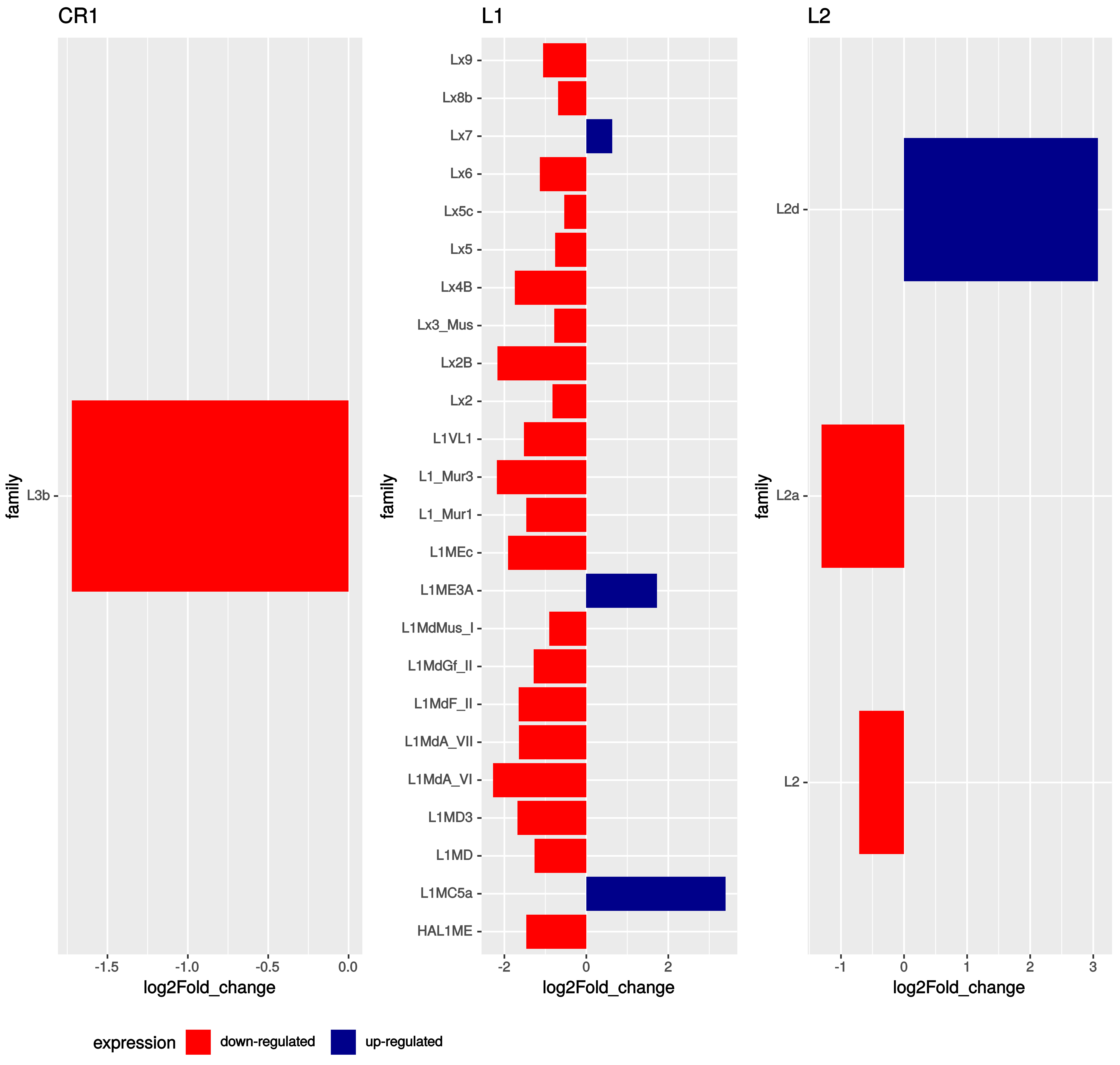


**Supplementary Fig. S7:** Log2fold change of differentially expressed families from the LINE retrotransposon class according to their superfamily. Red and blue bars indicate respectively down- and up-regulation.

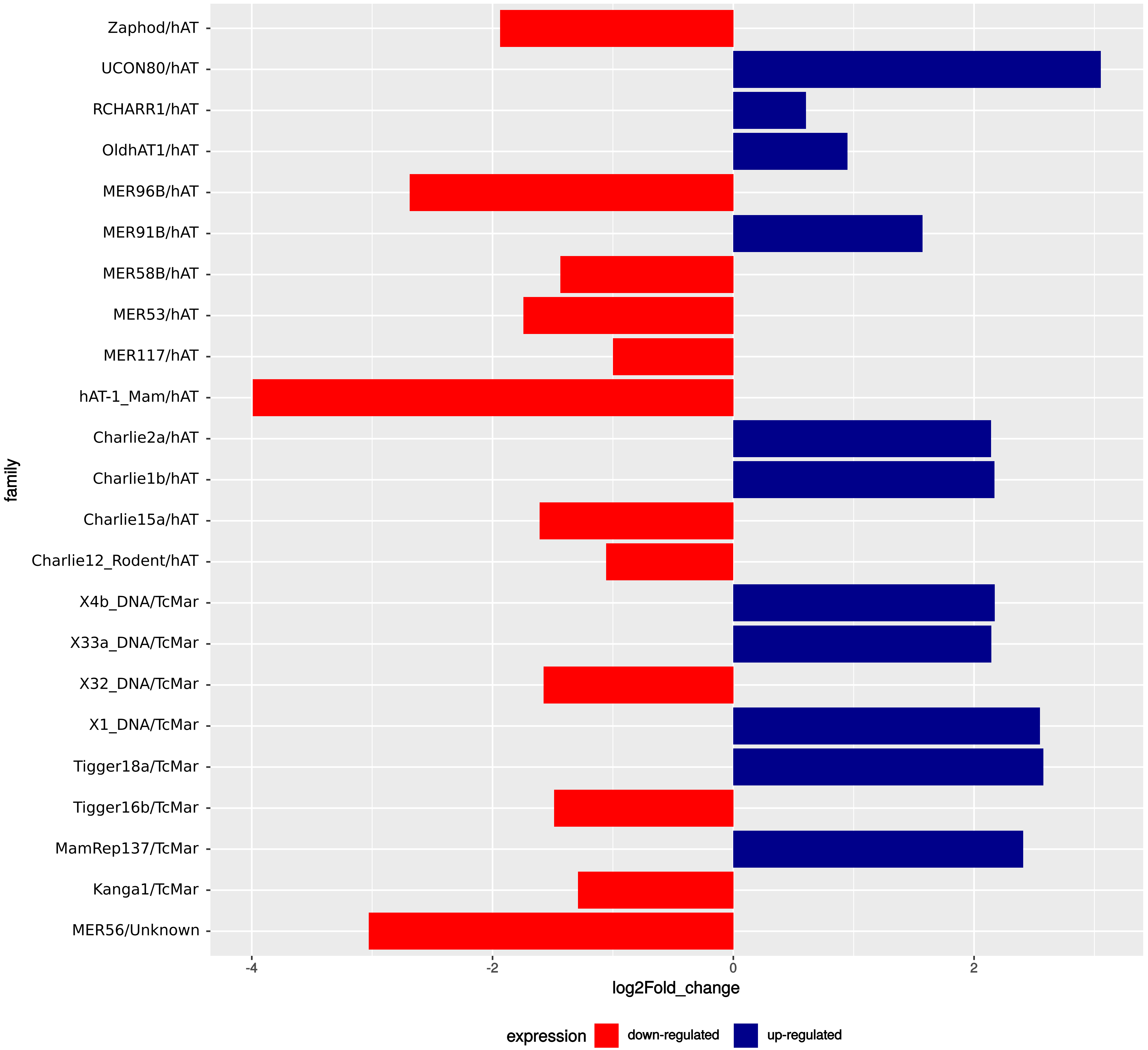


**Supplementary Fig. S8:** Log2fold change of differentially expressed families from the DNA transposon class Red and blue bars indicate respectively down- and up-regulation.

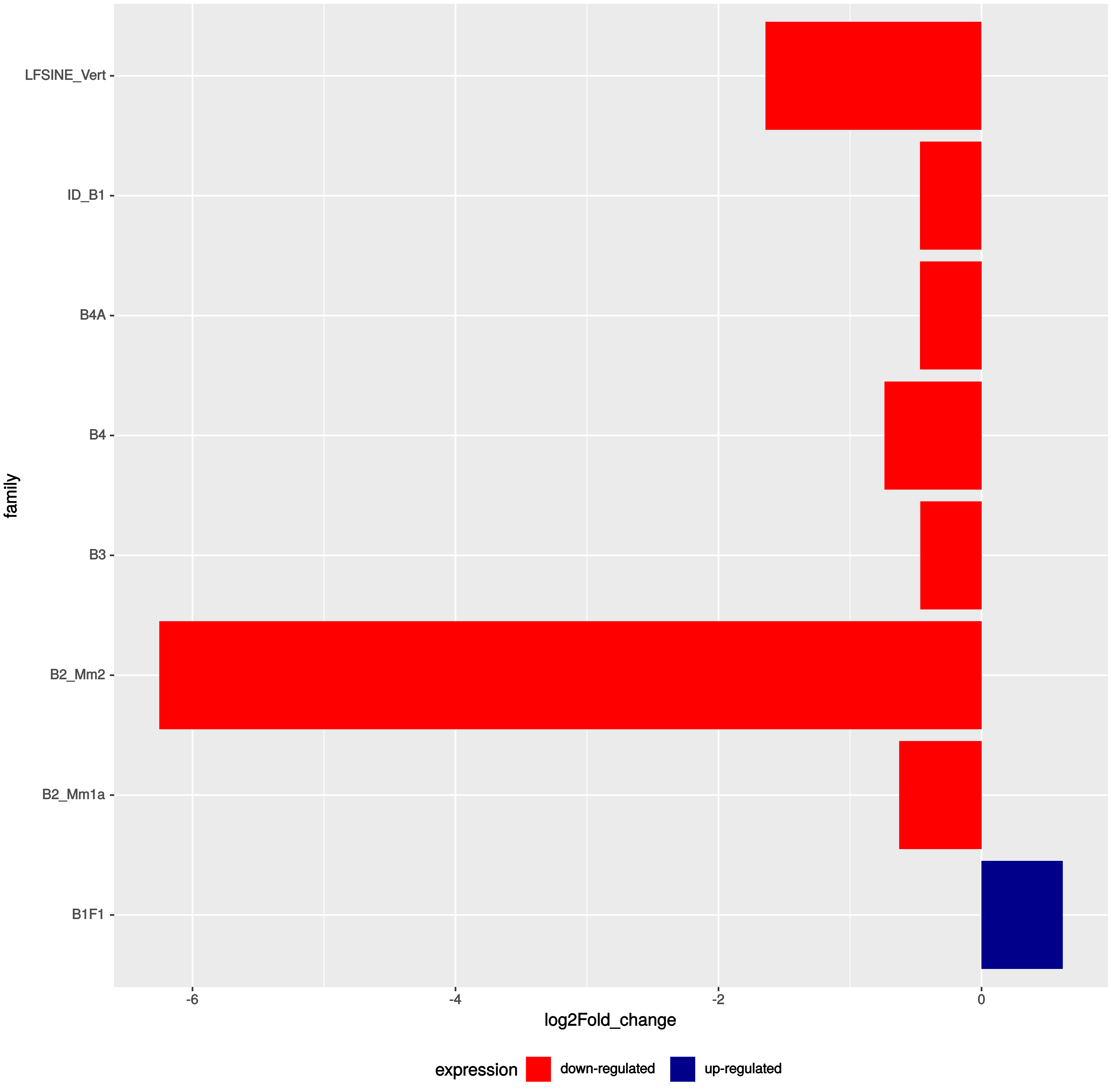


**Supplementary Fig. S9:** Log2fold change of differentially expressed families from the SINE non-LTR retrotransposon class. Red and blue bars indicate respectively down- and up-regulation.

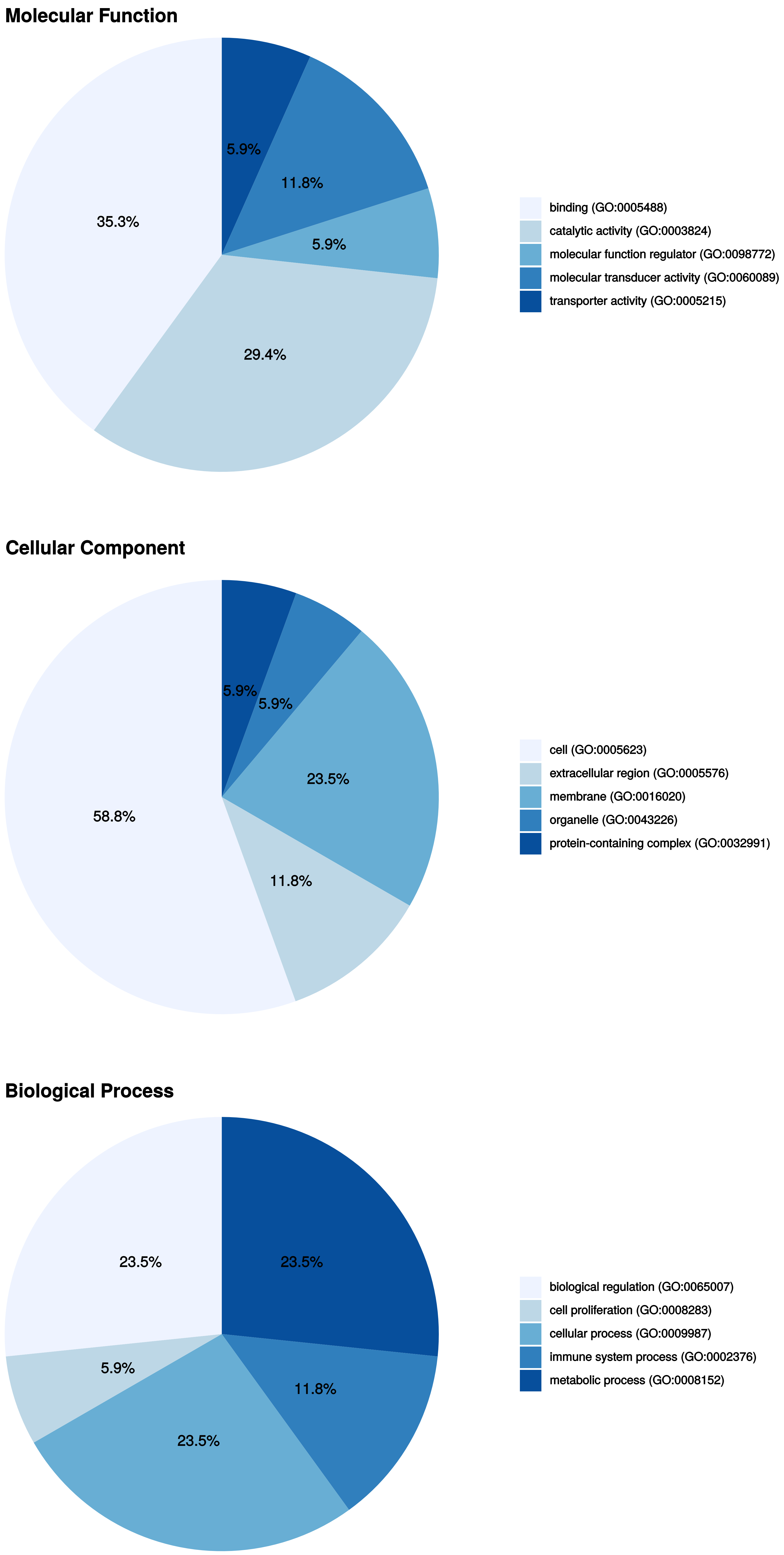


**Supplementary Fig. S10**: Function enrichment from GO analysis for chimerc transcript initiated in TEs that are differentially expressed between tumor and normal condition
